# A correlation analysis framework for localization-based super-resolution microscopy

**DOI:** 10.1101/125005

**Authors:** Joerg Schnitzbauer, Yina Wang, Matthew Bakalar, Baohui Chen, Tulip Nuwal, Shijie Zhao, Bo Huang

## Abstract

Super-resolution images reconstructed from single-molecule localizations can reveal cellular structures close to the macromolecular scale and are now being used routinely in many biomedical research applications. However, because of their coordinate-based representation, a widely applicable and unified analysis platform that can extract a quantitative description and biophysical parameters from these images is yet to be established. Here, we propose a conceptual framework for correlation analysis of coordinate-based super-resolution images using distance histograms. We demonstrate the application of this concept in multiple scenarios including image alignment, tracking of diffusing molecules, as well as for quantification of colocalization.

**Significance statement:** Correlation analysis is one of the most widely used image processing method. In the quantitative analysis of localization-based super-resolution images, there still lacks a generalized coordinate-based correlation analysis framework to take fully advantage of the super-resolution information. We show a coordinate-based correlation analysis framework for localization-based super-resolution microscopy. This framework is highly general and flexible in that it can be easily extended to model the effect of localization uncertainty, to the time domain and other distance definitions, enabling it to be adapted for a wide range of applications. Our work will greatly benefit the quantitative interpretation of super-resolution images and thus the biological application of super-resolution microscopy.

## Introduction

In the recent years, localization-based super-resolution microscopy has been demonstrated as a powerful technique to image beyond the diffraction limit and has produced numerous beautiful images of subcellular structures (1). The scientific community has now started to embrace it as a routine tool to answer actual biomedical questions. To make accurate conclusions about the biological system of interest, it is often required to quantitatively characterize the acquired images. While numerous analysis strategies have been well established for conventional fluorescence microscopy, these strategies do not apply directly to localization-based super-resolution images. The reason is that a conventional fluorescence image consists of pixels or voxels, a localization-based super-resolution image consists a collection of two- or three-dimensional coordinates, each associated with a localization uncertainty. Under this coordinate-based representation, many trivial operations on conventional images, such as thresholding and background subtraction, become challenging. A simple solution for adapting localization-based super-resolution images to established analysis routines is binning the coordinates on a pixel grid; however, this binning inevitably leads to a loss of precise localization information. On the other hand, certain operations complicated for conventional pixelated images, such as sub-pixel image translation, rotation and image deformation, become straightforward for coordinates. Therefore, it would be greatly beneficial to establish a generalized coordinate-based analysis framework.

We focus on image correlation, which is one of the most widely used image processing method. For localization-based super-resolution microscopy, correlation analysis or related methods (such as Ripley’s functions) have been used in measuring resolution (2), testing colocalization and clustering (3-5), image-based drift correction (6, 7), aligning super-resolution images of individual structures (8), although most of these cases still used spatial binning of the super-resolved coordinates.

Here, we present a coordinate-based correlation analysis framework (9, 10) for localization-based super-resolution microscopy. We mathematically showed that point-point distance distribution is equivalent to pixel-based correlation function. This point-point correlation function can easily model the effect of localization uncertainty. Moreover, this concept can be extended to the time domain and other distance definitions. We then demonstrated our framework in three applications of super-resolution microscopy: model-free image alignment and averaging for structural analysis, spatiotemporal correlation analysis for the mapping of molecule diffusion, and the quantification of spatial relationship between complex structures utilizing the generalized point-set distance definition.

## Results

### Correlation function between two coordinate-based images

In the pixel-based image representation, the translational cross-correlation function of two images A and B, *C_AB_*(*ξ, η*), is usually defined as the mean of pixel value products when shifting the two images by (ξ,η), normalized by the product of mean pixel values:

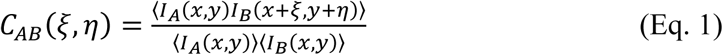

where *I_A_* and *I_B_* are brightness values at pixel (*x, y*). For localization-based super-resolution microscopy, images A and B are collections of *N_A_* and *N_B_* localization points, and thus they can be described as the sum of Dirac delta functions with each function representing one single-molecule localization point (11, 12). By replacing *I_A_* and *I_B_* of Eq. 1 with this coordinate-based definition, we can derive that the cross-correlation function between two coordinate-based images is (see Supplementary Note 1):

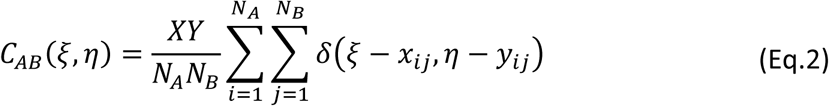

where *X* and *Y* are the image dimensions, and (*x_ij_*, *y_ij_*) is a vector from localization point *i* in image A to *j* in image B. Because the product of Dirac delta functions is non-zero only when their coordinates overlap, the correlation function of two coordinate-based images is yet another set of coordinates, located at the point-to-point vectors from coordinates in image A to coordinates in image B (Figure 1). In other words, the correlation function is a displacement map showing how image B has to be translated so that one point in image A overlaps with one point in image B. Therefore, we name *C_AB_*(*ξ, η*) the “point-point correlation function”.

**Figure 1.**
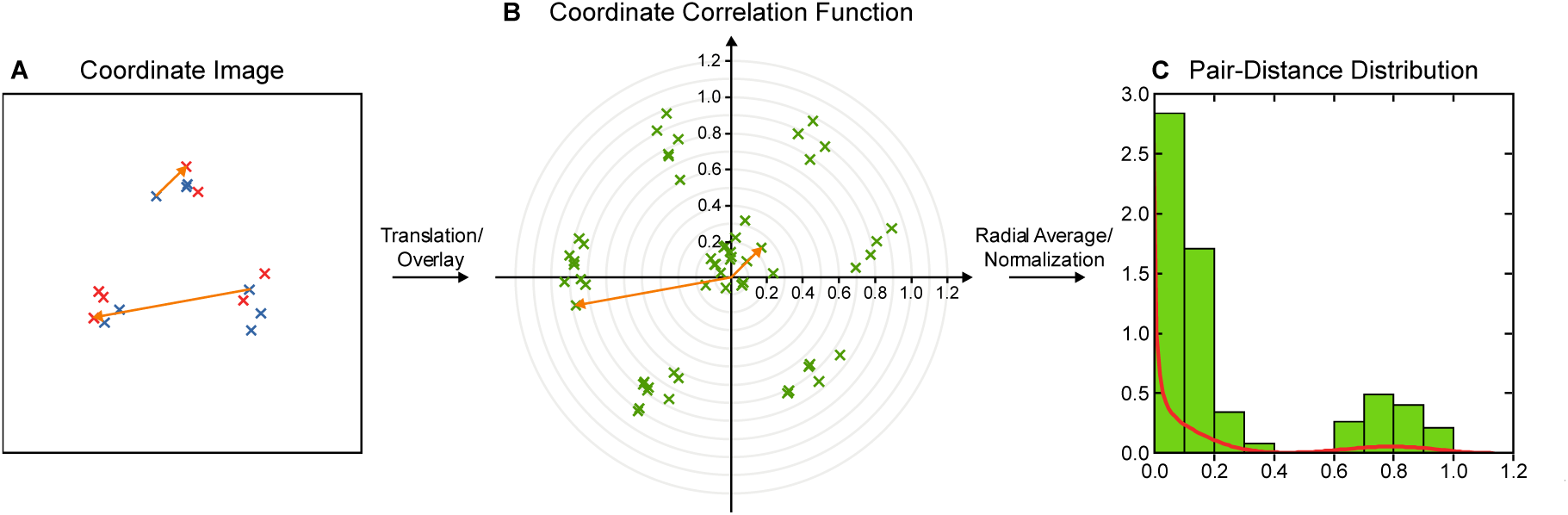
The coordinate-based correlation function and its relationship to the pair-distance distribution. The correlation function of coordinate-based images (red and blue) (A) is yet another set of coordinates (B). Each coordinate in correlation function (green) is located at the tip of a vector that connects one red localization with one blue. Two example vectors are shown in orange. (C) The normalized radial average of the correlation function shown in (B) is the pair-distance distribution, which can be either presented by binning (green) or as a kernel density (red) (see Supplementary Note 3).

In order to describe the effect that each localization point is associated with uncertainties in its coordinates, we replace the Delta functions with normal distributions. The two-dimensional point-point correlation can then be written as

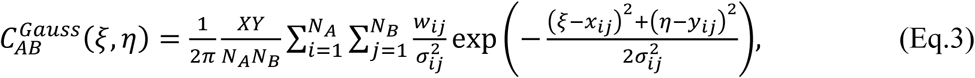

where 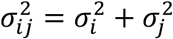 is the variance of the correlation vector connecting localization *i* with localization *j*, with precision variances 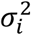 and 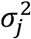, respectively. The weight factor, *w_ij_*, is used to weigh the correlation vectors, which is typically set to the inverse of local localization density around localization *i*.

When ignoring the directional information, a radially averaged 2D correlation function (see Supplementary Notes 2 and 4) forms a “pair-distance distribution”

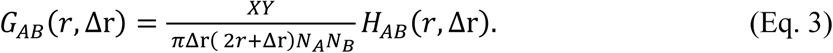

Here, *r* is the radial coordinate and Δ*r* is the radial bin size, and *H_AB_* (*r*, Δ*r*) is the number of pair-wise distances between A and B that fall into the [*r*, *r* + Δ*r*) range. Equivalent versions of *G_AB_*(*r*) were previously used to quantify clustering and co-localization in localization-based super-resolution microscopy under the term “pair correlation” (4) and “steady-state correlation” (5).

### Using image correlation for the alignment of super-resolution images

A straightforward application of image correction is in aligning and averaging multiple super-resolution images of a subcellular structure in order to gain signal-to-noise ratio. This very successful biological applications for super-resolution microscopy has allowed *in situ* dissection of molecular organizations for large protein complexes (8, 13-19). In most cases, though, image aligning and averaging relied on manual image stacking by hand (13), imposing a pre-defined structural model (16, 17), or pixel-binning the coordinate images (8, 18). Algorithms for model-free averaging of coordinate-based images emerged only recently (20).

We incorporate our coordinate-based definition of image correlation into an extensively used single-particle cryoEM reconstruction strategy (21). In this method, the sum of all images acted as the initial reference, and translational and rotational transformations were applied to individual images to maximize their correlation with the reference, then the reference was updated with the sum of transformed images. This procedure was iterated multiple times to result in a satisfactory alignment (Fig. 2A) (Online Methods). Compared to pixel-based implementations, the major advantage of using the coordinate-based image representations is that image transformation is straightforward and can be performed exactly without interpolation or loss of information.

**Figure 2.**
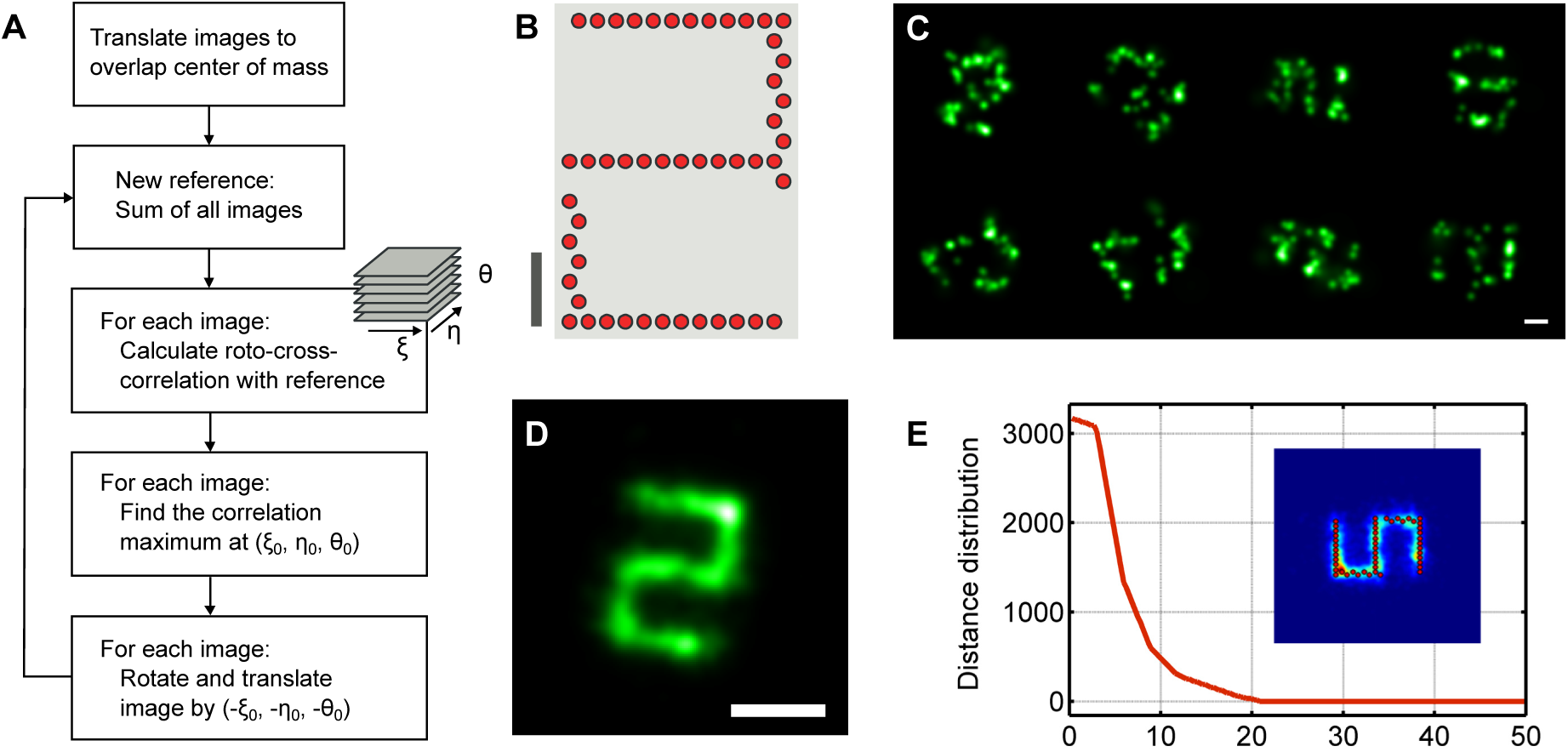
Structure refinement by translational and rotational image alignment. (A) Scheme of the alignment algorithm. (B) The DNA origami design. (C) Example images of individual structures reconstructed from DNA-PAINT localizations. (D) Sum of 133 individual DNA-PAINT images as in (C) after roto-translational alignment, revealing the underlying organization of DNA binding sites as the digit “5”. (E) pair-distance correlation between the ground truth and the aligned image, the insect shows the overlay of ground truth and the aligned image. Scale bars 50 nm.

As a demonstration, we applied this algorithm to DNA-PAINT images of a DNA origami structure (22) (Fig. 2B). We purposely used a very short imaging period (500 frames) so that each image is highly noisy because of the very limited number of localization points (Figs. 2C and S 1). The mean localization precision of this dataset is estimated to be 3.7 nm. By rotationally and translationally aligning and summing 133 images (no initial model was assumed), the underlying structure was precisely recovered (Fig. 2D). We further calculated the pair-distance distribution between the combined image and the ground truth from the origami design to quantify the alignment precision (Fig. 2E). The resulted correlation function has peak width of 4.9 nm, which is comparable to the mean localization precision, indicating that our coordinate-based model-free image alignment method has very high alignment precision.

### Frame-pair correlation for the mapping of molecule diffusion

Next, we show that by incorporating temporal information, pair-distance distribution analysis can be used to analyze the movement dynamics of biomolecules in living cells. By combining with photoswitching and single-molecule tracking, localization-based super-resolution microscopy has allowed a high density of target molecules to be labeled and followed over time (albeit with short trajectories) (23, 24), thus offering the opportunity to map the spatial heterogeneity of molecule diffusion and transport (24-26).

Tracking a moving molecule considers the displacement of fluorescent molecules from one camera frame to the next one to produce diffusion trajectories. Coincidently, a collection of all pair-wise displacements is exactly the cross-correlation function of molecule localizations in these two camera frames. Therefore, we define the frame-pair distance distribution FPD(*ξ*, *η*) as the average displacement map of molecules localized in two consecutive frames (showing in the radially averaged form):

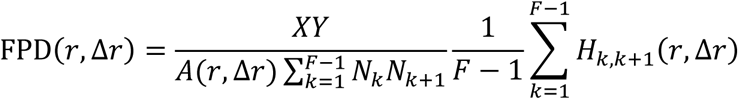

Here, *F* is the total number of frames, *k* is the frame number indicating single data acquisition time point and *H*_*k, k* + 1_(*r*, Δ*r*) is a histogram of distances between localizations in frame *k* with frame *k* + 1. This FPD is analogous to Image Correlation Spectroscopy (27), Particle Image Correlation Spectroscopy (PICS) (11), and the localization-specific Spatio-Temporal Image Correlation Spectroscopy (STICS) (28).

FPD describe the ensemble molecule diffusion activity within the area of analysis. For two-dimensional Brownian diffusion, the resulted FPD distribution is a Gaussian peak centered at zero, with a standard deviation representing the mean displacement (MD) per frame. MD is determined by the diffusion coefficient, *D*, the delay between two exposures, Δ*t*, and the localization precision, *p*:

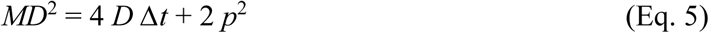

FPD is particularly useful for the spatial mapping of diffusion behavior because such measurement require a high density of tracked molecules. At this density, single-molecule tracking, becomes difficult because collision of molecules, together with their constant activation, blinking and bleaching lead to incorrected linked trajectories (29), whereas FPD is unaffected because these effects only contribute to a flat baseline that can be easily handled.

To demonstrate FPD-based diffusion analysis, we took STORM images of *Drosophila* S2 cell membranes stained with a photoswitchable membrane dye DiD-C_18_ (25) (Fig. 3A). Because DiD is a small molecule that has a high diffusion speed in the membrane, we took a stroboscopic illumination scheme to reduce the motion-blur from fluorophore diffusion within the exposure time (30). While the camera exposure time was 8.3 ms (~121 Hz frame per second), we turned on the excitation laser only for 1/10 of the frame duration (0.83 ms). Moreover, by varying the time point of strobing within a frame, we were able to access sub-frame temporal resolution. Specifically, we turned on the laser at 8/10 of the frame duration for even frames and at the beginning of a frame for odd frames (Fig. 3B). As a result, the effective time lag from even to odd frames was only 0.2 frames (1.7 ms) and 1.8 frames (15 ms) from odd to even frames. We then computed the frame-pair distance distribution for these two different sets of frame pairs (Fig. 3C). An effective diffusion coefficient, *D*_eff_, can be calculated for either of the two curves by fitting the peak width assuming normal Brownian diffusion and considering a localization precision *p* of 17 nm (calculated from the photon number and background values). The value of *D*_eff_, at ~ 0.5 μm^2^ / s for both lag times, is consistent with previous measurement of DiI/DiD diffusion in cell plasma membrane (31). Benefited from the high localization density, we were able to produce a spatial map of *D*_eff_ by binning the STORM image and fitting the frame-pair distance distribution of each spatial bin. The *D*_eff_ maps appear rather homogenous across the cell plasma membrane (Figs. 3E & F).

**Figure 3:**
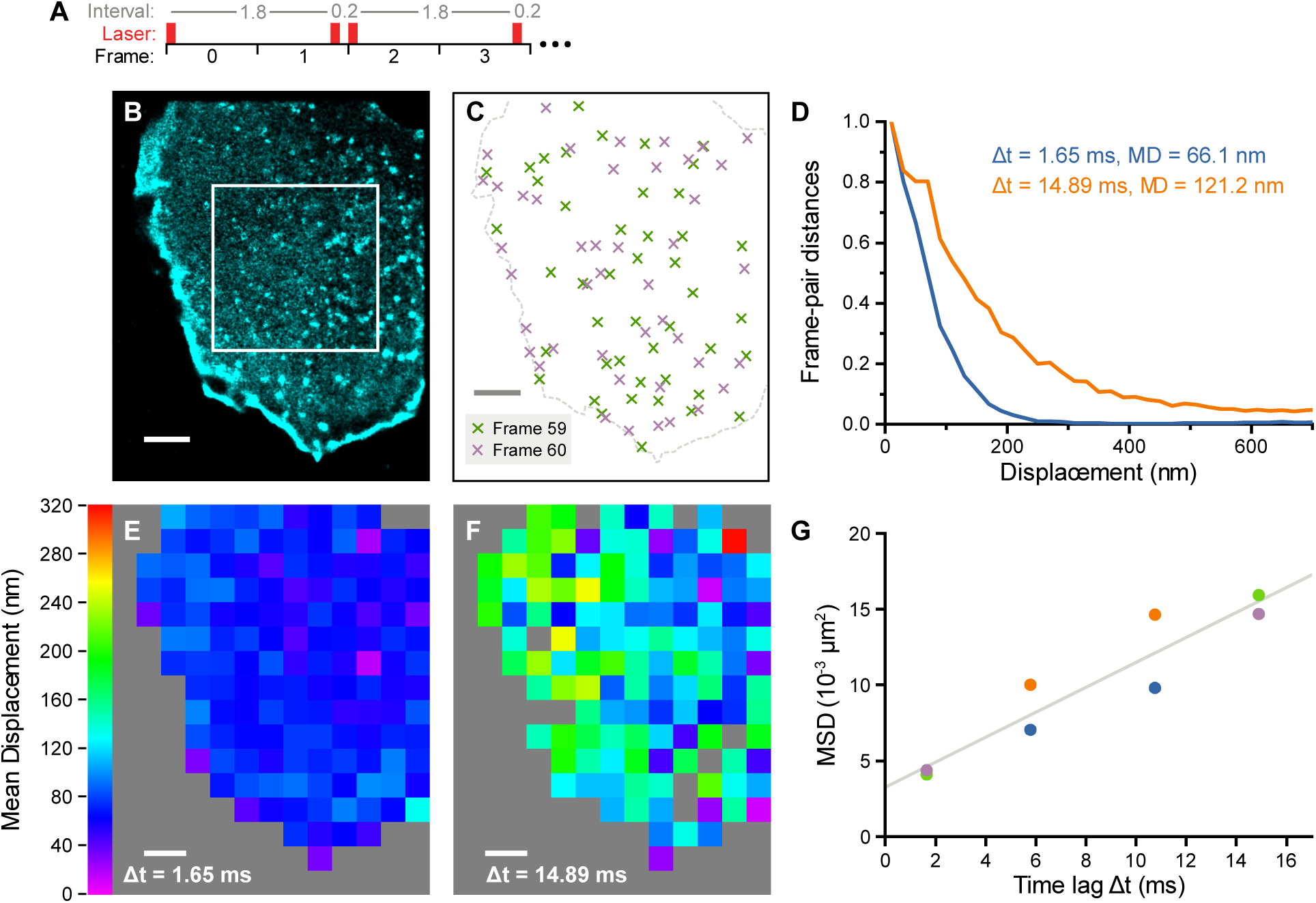
Diffusion measurement by frame-pair correlation analysis of coordinate-based super-resolution images. (A) Strobing sequence of the excitation laser synchronized to the camera frames. (B) Super-resolution image of S2 cell membrane stained by DiD (left) and localization points in two consecutive frames, illustrating the high molecule density. (C) Frame-pair distance distribution of localizations in boxed area of (A) for two time lags, normalized to the interval [0, 1]. (D) Maps of effective DiD diffusion coefficient calculated for the two time lags. (G) Mean Square Displacement (MSD) obtained from fitting the frame-pair correlation function using different laser shutter intervals. Each color represents a different cell. Scale bars 1000 nm.

With additional sets of measurements at lag times of 5.8 and 10.8 ms, we were able to construct a mean squared displacement curve at the time range of 1.7 to 14.9 ms (Fig. 3G). Linear fitting indicates that DiD undergoes largely simple Brownian diffusion at this time scale, which is consistent with previous STED-FCS studies of simple lipids (32). In fact, our variable-time strobe method probes into a comparable temporal (1.7 ms) and spatial (41 nm FWHM for localization precision) scale as STED-FCS (33), and it has the similar capability to resolve heterogeneous diffusion behavior by fitting the frame-pair correlation curve with a multi-species diffusion model. Therefore, it offers a complementary method to investigate lipid diffusion and membrane heterogeneity at high spatiotemporal resolution, with STED-FCS potentially offering a higher temporal resolution, and variable-time strobe proving an easier way for spatial mapping.

### Point-to-set correlation function for structural and colocalization analysis

Finally, exploiting the equivalence between the distance histogram and the correlation function of coordinate-based super-resolution images, alternative definition of distances can be incorporated in our framework for quantifying the spatial relationship between more complicated structures. Specifically, by considering the distance between individual point A and all points in B as a set, ***B***, we used the histogram of these point-set distances to define a point-set correlation function. Mathematically, the distance between a point and a set of points is defined as the distance between this point and its nearest neighbor in the set. We calculate the histogram of this distance for all A points, *H_AB_*(*r*, Δ*r*), and then normalize it by the image area occupied by each bin of the histogram, *A_B_*(*r*, Δ*r*), and the average localization density in the whole field of view (Fig. 4A)

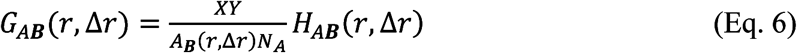

This correlation function coverts the point sets in the original image into a numerical function, which allows us to correct the contributions from channel cross-talk arithmetically (see Supplementary Note 5).

**Figure 4.**
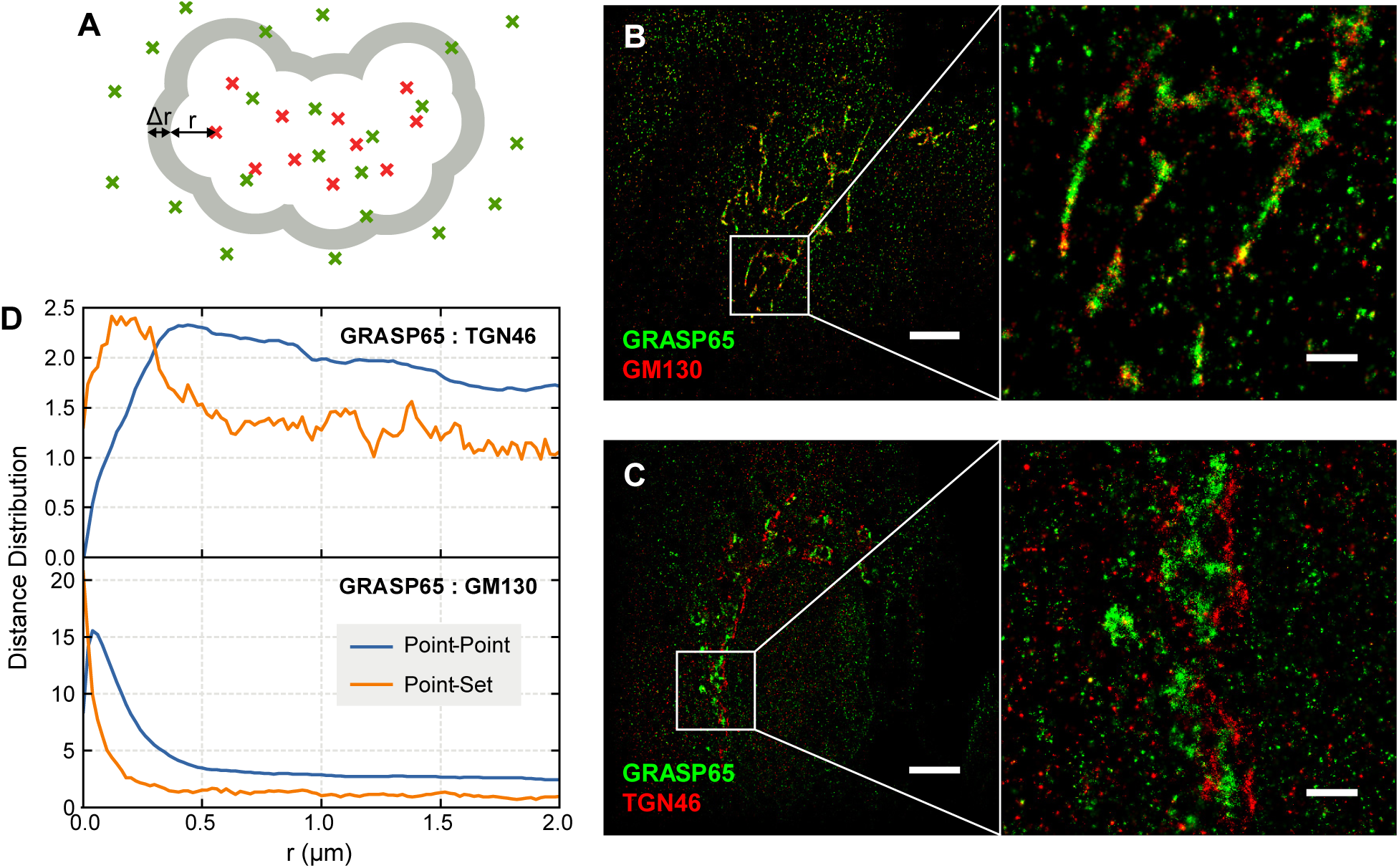
Co-localization and distance characterization using the point-set correlation function. (A) Schematic of the point-set correlation function. Red and green crosses represent the set and point localizations, respectively. The point-set correlation function is the histogram of the number of green localizations (points) with a distance between *r* and *r* + Δ*r* to any red localization (set), normalized by the area from which the corresponding distance vectors could originate (grey), and by the average “point” density. (B and C) Two-color STORM images of GRASP65 (green) with (B) cis-Golgi marker GM130 and (C) trans-Golgi marker TGN46. Scale bars 5 μm for the overviews and 1 μm for the zoomed-in panels. (D) Point-point and point-set cross-correlation functions of (B) and (C).

To illustrate the utility of the point-set correction concept, we analyzed the spatial relationship between the cis-Golgi protein GRASP65 and either the cis-Golgi protein GM130 (Fig. 4B) or the trans-Golgi protein TGN46 (Fig. 4C) (34). We devised an algorithm to automatically identify Golgi ribbons in either the GM130 or TGN46 channel, calculated the point-set and point-point correlation functions between GRASP65 and Golgi ribbons, and then subtracted the crosstalk contribution from the correlation functions (Fig. 4D). For GRASP65 and GM130, which was shown biochemically to interact with each other (35), the point-set correlation displayed a sharp peak at zero distance, indicating strong colocalization. On the other hand, the point-point correlation peak was broader (reflecting the width of the Golgi ribbons) and contained a dip at zero distance. This dip may be due to spatial exclusion of antibodies, differential accessibility of the two epitopes, or imprecise cross-talk subtraction. For GRASP65 and TGN46 which should exist in parallel, non-overlapping structures, the point-set correlation displayed a peak at around 200- 300 nm, reflecting the distance between cis- and trans-Golgi ribbons. In contrast, the point-point correlation was completely smeared by the length of the Golgi ribbons, making it much less informative than the point-set correlation in characterizing the spatial relationship between these two parts of Golgi.

## Conclusion

In summary, we have demonstrated the utility of our coordinate-based correlation analysis framework in quantitative interpretation of localization-based super-resolution microscopy data in a number of cases: image alignment, tracking of molecular diffusion and quantification of colocalization. We have also shown the generality and flexibility of this framework in expanding into the time domain (frame-pair correlation) and adoption alternative definitions of distance (point-set correlation). Although we described our framework and algorithms in two-dimensional representation, they can be easily extended to 3D super-resolution microscopy. The resulted correlation function will be a set of points at the tip of three-dimensional distance vectors between two localizations. Even though in most cases, the localization precision in the axial direction is different from that in the lateral direction, this anisotropy can be handled by transforming the uncertainty cloud together with image transformation. We expect that the analysis methods described here, as well as model-free super-resolution image alignment and fast diffusion analysis by strobe illumination with variable-timing, will be broadly useful in practical applications of super-resolution microscopy in various biological systems. Finally, we envision that in many cases, the most efficient algorithm will be a hybrid of coordinate- and pixel-based approaches (Supplementary Note 6).

## Materials and methods

### Roto-translational Image Alignment

The experimental conditions for this dataset were described previously (22). A set of 133 images of the origami structure was picked by hand (Supplementary Figure 1). The alignment procedure was performed as follows. First, coordinates of individual images are shifted so that their center of mass overlaps. Then, several iterations of the following procedure were applied: (1) for each individual image, find the translation and rotation that maximizes the cross-correlation of this image with the sum of all other images. For that matter, a brute force maximizing algorithm was used that samples the cross-correlation function at discrete steps of the rotation angle and the two translational dimensions. For a given rotation, the translational cross-correlation was evaluated as the sum of Gaussians, centered at the tip of pairwise distance vectors with a width of the combined localization uncertainty (theoretically estimated as in (36)). Note that this brute force approach is simple, yet highly inefficient. However, the algorithm could easily be improved by using a typical optimization problem solver (even the Jacobian can be supplied as the gradient of the sum of Gaussians). (2) Apply the correlation-maximizing rotation and translation to each images, so that a new and improved image is formed when taking the sum of all images. For the alignment shown in this paper, ten iterations of this procedure was applied with a 5 nm translational grid (+/- 25 nm around the center of mass in each dimension) and 5° rotational step size (covering the complete 360°). The resulted aligned images were further optimized by three iterations with a 2 nm translational grid (+/- 10 nm around the center of mass) and a 2° rotational step size (again covering 360°). We found that image alignment is strongly dominated by local clusters comprising a relatively high number of localizations. Consequently, the algorithm often aligned the brightest clusters of the individual images to overlap, while almost disregarding the rest of the structure. This feature likely arises for two reasons: (a) images obtained by localization microscopy tend to appear artificially clustered due to the highly stochastic blinking process; and (b) the cross-correlation function has a squared dependency on image intensity. Hence, to avoid overestimation of artificial clustering in this algorithm, the distance vectors were weighted by local density of the localization in the individual image. Specifically, the Gaussian volume for a distance vector was set to 1/*N_local_*, where *N_local_* is the number of localizations that are closer than twice the localization precision of the vector origin (in the individual image).

### Single-Molecule Localization Microscopy of DiD in S2 Cells

Wild-type S2 cells were cultured in Sf-900 II serum free medium (Gibco) and plated onto 35 mm glass bottom dishes no. 1.5 cover glass (MatTek) for fluorescence imaging. To facilitate a spread out cell morphology, the coverglass was coated with 0.1 mg/ml concanavalin A for 0.5 h before cell plating. The cells were then washed (3x, PBS), fixed (4% Paraformaldehyde, 10 minutes), incubated in DiD (1 μM, 30 seconds) and washed again (3x PBS). For images, cells were mounted in imaging buffer (PBS, 1 M mercaptoethylamine, pH 8.5, 50% glucose in MilliQ water and oxygen scavenging solution (10 mg of glucose oxidase, 25 μl of catalase in 100 μl of PBS) in the ratio of 80:10:10:1). Single molecule localization data acquired on a microscope described before (37) with 121 Hz camera frame rate, 128 by 128 pixel field of view and EM gain 100. For DiD excitation the 642 nm laser power was ~74 mW measured at the back port of the microscope. The laser was shuttered in synchronization with the camera frames as shown in Figure 3A.

### STORM and Image Analysis of Golgi Proteins

RPE cells were seeded overnight in a Lab-Tek II eight-well chambered no. 1.5 cover glass (Nunc), washed twice with warm PBS and fixed with warm 4% PFA for 15 minutes at 37°C. Then, cells were washed again (3x, PBS) and blocked with 3% BSA and 0.5% Triton-X 100 in PBS for 15 min. Primary antibody (Rabbit Anti-GRASP65, ab30315, Abcam; Sheep Anti-TGN46, AHP500GT, AbD Serotec; Goat Anti-GM130, sc-16268, Santa Cruz; 1-5 μg/ml) staining was performed overnight in 3% BSA at 4°C, followed by washing (3x, PBS) and secondary antibody staining (1-5 μg/ml) in 3% BSA for one hour. Cells were washed again (3x, PBS) and post-fixed with 3% PFA + 0.1% Gluteraldehyde in PBS before final washing (3x, PBS). Secondary antibodies were labeled with a mixture of two fluorophores (either Alexa Fluor 405 and Alexa Fluor 647, or Cy3 and Alexa Fluor 647) for activator-reporter type two-color STORM imaging (6). STORM acquisition was performed on a microscope described before (37). For display, non-specific blinking was removed from the STORM images (13). The point-set distance distribution was calculated as follows. First, background was removed from the reference channel (GM130 or TGN46). For that matter, the local density of each localization in the respective channel was calculated by counting localizations of the same color within a radius of 300 nm. Localizations with a local density lower than a manually applied threshold were subsequently removed from the image. The remaining molecules are defined as the “set” for the following computation. For the point-set distance distribution, the GRASP65 density was calculated as a function of distance *r* to the reference set (GM130 or TGN46). To facilitate the computation of the area of each distance shell around the reference set, the calculation was performed based on pixelated STORM images (20 nm pixel size) where the pixel value is equal to the number of localizations in that pixel. The GRASP65 pixel image was subtracted by a pixel image from non-specific blinking events (which was divided by the ratio of specific to non-specific frames beforehand) (6, 13), allowing negative values to preserve statistical effects. The image of the reference set was convolved with a centered logical disk of radius *r* (each pixel is either unity if its distance to the image center is less than *r* and zero otherwise). The resulted convolution is a logical map of pixels that are closer than *r* to the reference set. Based on logical maps for a range of distances *r*, the cumulative point-set distance histogram was calculated by summing up all pixels in the GRASP65 image where the logical map is equal to or larger than unity. Besides the localization histogram, the corresponding cumulative area was calculated by summing up all non-zero pixels in the logical map. The point-set distance distribution was then calculated by taking the discrete difference of the cumulative localization histogram and dividing the result by the discrete difference of the cumulative area.

## Acknowledgement

We thank Francis Brodsky and Yanzhuang Wang for the help with Golgi staining and imaging. Ralf Jungmann and Peng Yin generously provided the DNA-PAINT data. This project is supported by the NIH Director’s New Innovator Award (DP2OD008479), J.S. thankfully acknowledges support from a Boehringer Ingelheim Fonds Ph.D. fellowship, and S.Z. thank the School of Life Sciences, Peking University for the internship support. B.H. is a Chan Zuckerberg Biohub investigator.

## Author contributions

B.H. and J.S. conceived and designed the project. S.Z., M.B., T.N. and J.S. performed the DiD experiments. B.C. and J.S. performed the Golgi experiments. J.S. and B.H. analyzed the data. J.S. Y.W. and B.H. wrote the manuscript.

## Supplementary Note 1: The correlation function of coordinate-based images

The spatial cross-correlation of two images can be defined as

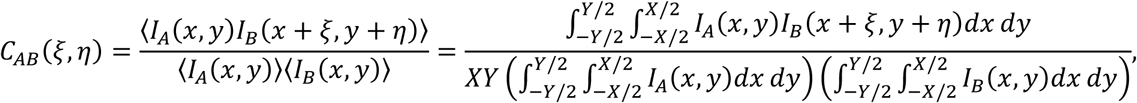

where *I_A_* and *I_B_* are images with coordinates *x*, *y*, and *ξ*, *η* are the correlation coordinates (Digman et al., 2005). *X* and *Y* are the image dimensions. For super-resolution localization microscopy, we define images A and B as the sum of two-dimensional Dirac delta functions:

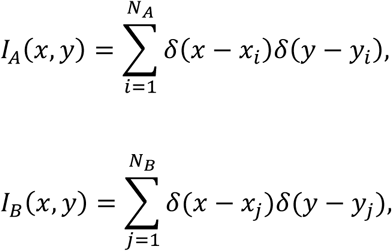

where *N_A_*, *N_B_* are the total number of localizations in image A and B and (*x_i/j_*, *y_i/j_*) are the coordinates of a localization *i* in image A or localization *j* in image B. Therefore, we can write the correlation function for coordinate based images as

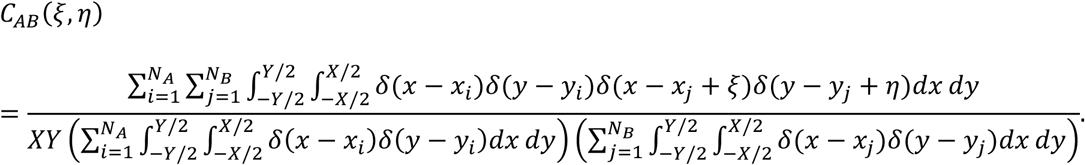

We now describe the coordinates of image B in terms of coordinates of image A:

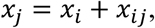

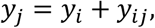

where (*x_ij_, Y_ij_*) is a vector from localization *i* in image A to localization *j* in image B. Substituting these and evaluating the integrals in the denominator result in

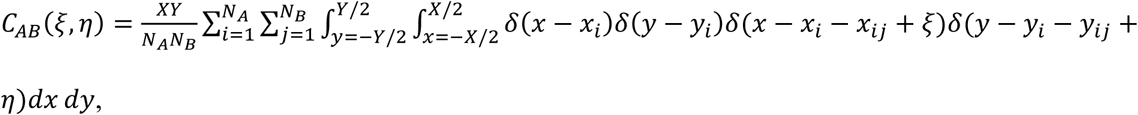

which leads to

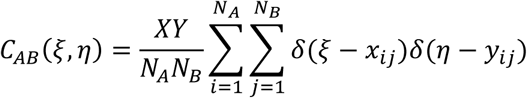

after evaluating the remaining integrals. Hence, the correlation function of a coordinate based image is the sum of delta functions located at the pair-wise distance vectors of coordinates in image A to coordinates in image B. We name this function therefore the “point-point correlation”.

## Supplementary Note 2: The radial average of the coordinate correlation function

In many situations, the two dimensional point-point correlation function is radially symmetric. Therefore the dimensionality can be reduced by defining its radial average as the “pair-distance distribution”, which resembles a histogram of pair-wise localization distances. With the radial and angular coordinates *ρ* and *φ*, defined by

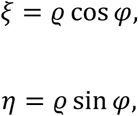

we can re-write the correlation function in polar coordinates:

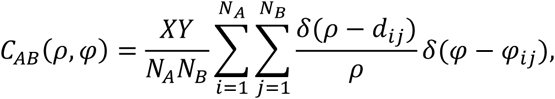

where *d_ij_* is the radial component of a correlation vector (the absolute distance between two coordinates) and *φ_ij_* is the angular component. Finally, the radial average is:

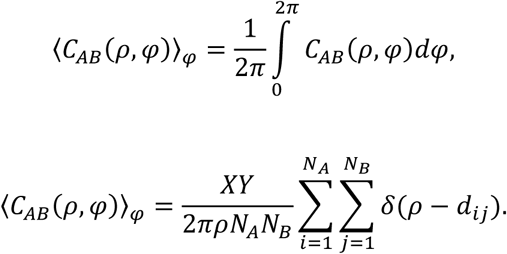

Hence, the one-dimensional pair-distance distribution is a sum of Delta functions located at the absolute distances between localization pairs in image A and B, with a normalization factor resulting from the image and radial averaging.

## Supplementary Note 3: Including localization uncertainty and accessing the correlation value via kernel density estimation

For single molecule localizations, each coordinate is assigned a specific localization uncertainty. Hence, the point-point correlation function can be extended by representing each correlation coordinate with a probability distribution instead of a Dirac peak. This probability distribution can be approximated by a 2D multivariate normal distribution with a variance being the summed uncertainty variance of the respective localization pair. Hence, the two-dimensional point-point correlation can be written as

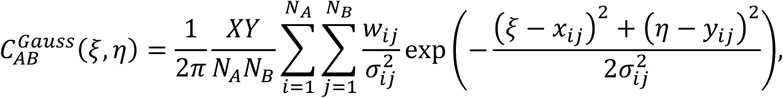

where 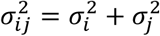 is the variance of the correlation vector connecting localization *i* with localization *j*, with precision variances 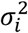and 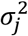, respectively. *w_ij_* can be used to weigh the correlation vectors (typically set to unity). For some applications (for example the here reported image alignment), it might be useful to set *w_ij_* to the inverse of local localization density around, for example, localization *i*. This might help reducing the overestimation of high density clusters which are a common artifact in localization microscopy.

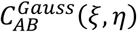 is functionally a two-dimensional kernel density estimator where the samples are the pair-wise distance vectors and the width of each sample’s kernel is determined by the pair of localization uncertainties. With additional knowledge about the kernel distribution, a more accurate representation than the normal distribution might be plugged in instead.

Analogously, the one-dimensional pair-distance distribution can be extended by approximating the distance probability with a normal distribution:

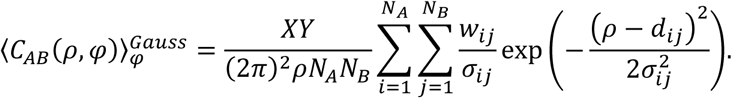

The function values of both 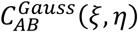 and 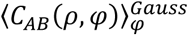 are well accessible for analytical or numerical methods, for example to find the correlation maximum with an optimization algorithm. Depending on the specific implementation, it might be useful to supply the Jacobian of the correlation function to the optimization algorithm, *e.g.* for the two-dimensional case:

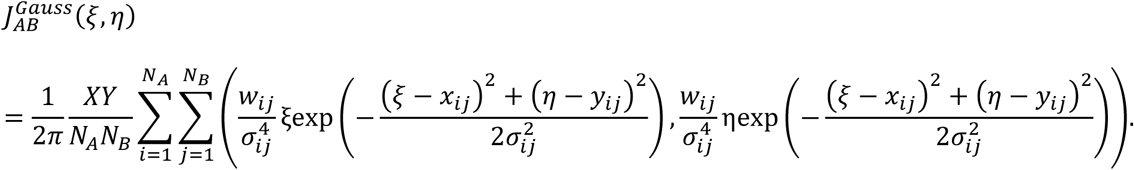

## Supplementary Note 4: Accessing the correlation value by binning

As an alternative to the kernel density estimation as described in Supplementary Note 3, the value of the radially averaged correlation function can be accessed by radial binning. The result is a normalized pair-distance histogram. With a bin size of Δ*r* and a radial coordinate *r*, the binned pair-distance distribution *G_AB_*(*r*, Δ*r*) is

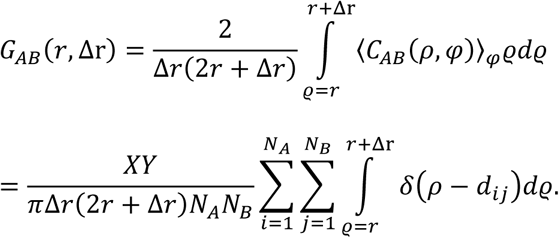

And finally

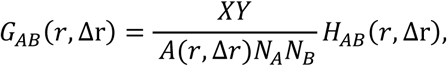

where *A*(*r*, Δ*r*) = *π*Δr(2*r* + Δ*r*) is the area of a shell with inner radius *r* and outer radius *r* + Δ*r*, and

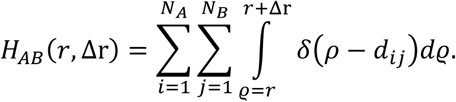

The spatial integration in *H_AB_*(*r*, Δ*r*) yields unity for pair-wise distances that are between *r* and *r* + Δ*r*, and zero otherwise. Therefore, *H_AB_*(*r*, Δ*r*) is a histogram of pair-wise absolute distances between each localization in image A to each localization in image B, with bin size Δ*r*. *G_AB_*(*r*, Δ*r*) is equivalent to the previously used pair-correlation (Sengupta et al., 2011). Several normalization factors of *H_AB_*(*r*, Δ*r*) give *G_AB_*(*r*, Δ*r*) a comprehensive meaning:

- 1/*N_A_*: Together with the summation over all image A localizations, this turns the function into the average histogram of the distance between a single localization in image A and all localizations in image B.
- 1/*A*(*r*, Δ*r*): By dividing the histogram value of each bin by the area of the corresponding shell, the function reports the density of image B localizations, instead of an absolute number.
- *XY/N_B_*: This number is equal to the inverse overall density of image B localizations in the whole field of view. Hence, the function value is relative to a hypothetical Poisson distribution of image B localizations.

In summary, *G_AB_*(*r*, Δ*r*) reports the average density of image B localizations at distance *r* from image A localizations relative to a hypothetical random distribution of image B localizations. More qualitatively, *G_AB_*(*r*, Δ*r*) reports how enriched localizations in image B are at a distance *r* to a localization in image A.

## Supplementary Note 5: Cross-talk subtraction in correlation functions

When the cross-talk between two or more colors in a localization-based image can be estimated, it is possible to correct the cross-correlation function between the respective colors. For example, STORM data in an activator-reporter multi-color schemes suffers from significant cross-talk between the colors due to the channel-unspecific blinking of the reporter dye (Bates et al., 2007). To subtract the cross-talk for visualization, random removal of localization points based on local density in the unspecific channel has been successful (Dani et al., 2010). For the case of quantitative analysis however, this approach is not suitable, due to the potential of underestimating cross-talk when no more localizations are present to be removed. Therefore, cross-talk must be considered on the level of the correlation function. The corrected correlation function is determined from corrected coordinate images 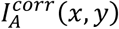 and 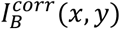:

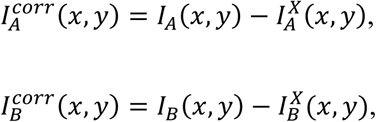

where 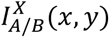 is the estimated cross-talk in image A or B, respectively. The cross-correlation is then

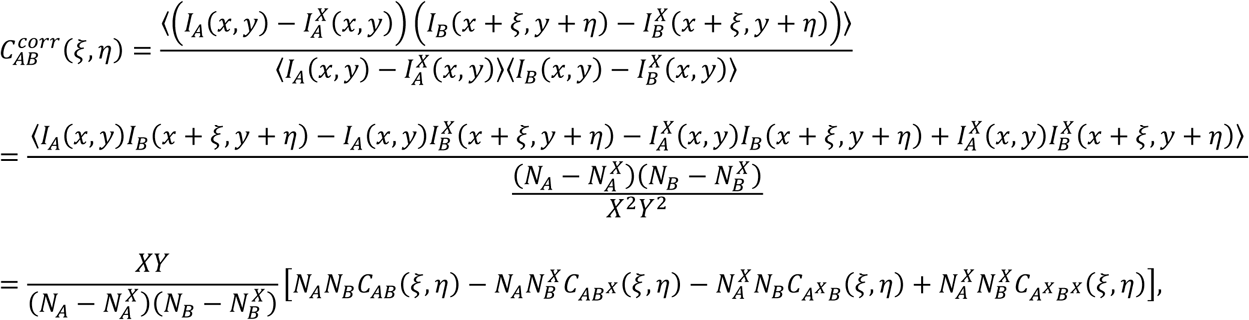

where 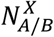 is the estimated number of cross-talk localizations appearing in image A or B, respectively. The X superscript in the correlation identifier corresponds to the expected image of cross-talk localization in respective image, so for example *C_A_x_B_*(*ξ, η*) is the cross-correlation of 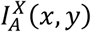 and *I_B_*(*x, y*).

Analogously, the corrected radial average (pair-distance distribution) is:

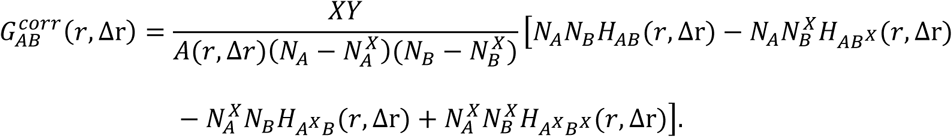

For the activator-reporter scheme of STORM, the estimated cross-talk in the two images is identical, hence the above derivation can be simplified with 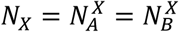 and 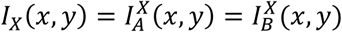.

## Supplementary Note 6: Comparison of coordinate-based and pixel-based correlation computation speed

We mathematically demonstrated that coordinate-based and pixel-based correlation function are fundamentally equivalent, which means all analysis described here can be implemented in either representations, and it has been previously shown that with a binning pixel size no larger than one fourth of the desired resolution, the pixilation effect on correlation calculation can be neglected (Nieuwenhuizen et al., 2013). Practically, computation time is one of the major factors affecting the choice of pixel versus coordinate image representation because of their different time complexity. Coordinate-based computation time scales with *N_L_*^2^, where *N_L_* is the number of localization points. In contrast, if Fast Fourier Transform (FFT) can be used, pixel-based computation time scales with *N*_P_log*N*_P_, where *N*_P_ is the number of pixels.

As an illustration, we computed the auto-correlation and the frame-pair correlation of a STORM image using both methods. We pixelated a super-resolution STORM image with 1,969,990 localizations in 49,995 frames on a 4096 by 4096 grid for FFT computation (bin size 8.9 nm). Both auto-correlation and frame-pair correlation were radially averaged. The radial average was evaluated over a range half the size of the STORM image with 2048 bins, except for the auto-correlation function computed by brute force looping, which was only evaluated in the first 512 bins (until *r*= 4576 nm) to avoid unreasonably long computation times. The two-dimensional correlation function *C*(*ξ*, *η*) of the pixelated images *I_A_*(*x*, *y*) and *I_B_*(*x*, *y*) was evaluated via FFTs by

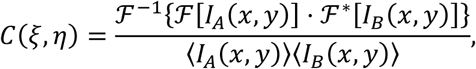

where 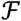 denotes the Fourier Transform and * complex conjugation. In case of the auto-correlation, the FFT method was about six orders of magnitude faster (3.6 ± 1 seconds (*n* = 10) versus ~ 111 ± 2 hours (*n* = 4)). In contrast, the computation of the frame-pair correlation was ~246 times faster with the looping method (14.6 ± 0.7 seconds (*n* = 10) versus 60 ± 4 minutes by the FFT algorithm (*n* = 5)). It is also intuitive that extending the calculation into three dimensions will substantially increase the computation time and memory demand for pixel- (voxel-) based calculation, but coordinate-based calculation will not be affected as long as the number of localization points remains the same. We envision that in many cases, the most efficient algorithm can be realized by a hybrid of coordinate- and pixel-based image representations.

## Supplementary Figure

**Supplementary Figure 1:**
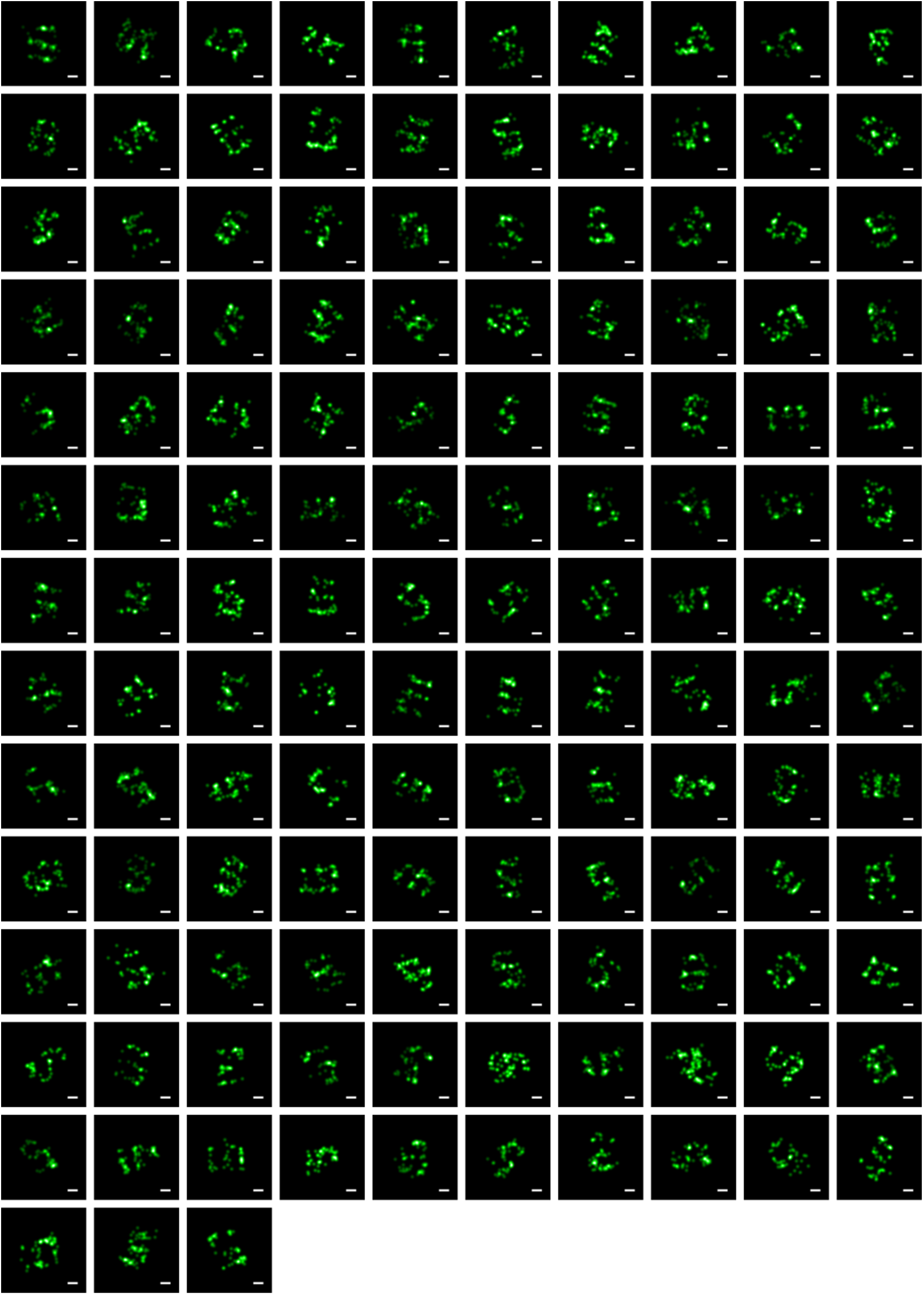
133 individual images that were aligned for the average image in Figure 4B. Scale bars 25 nm.

